# Phylogenetic profiling in eukaryotes: The effect of species, orthologous group, and interactome selection on protein interaction prediction

**DOI:** 10.1101/2021.05.05.442724

**Authors:** Eva S. Deutekom, Teunis J.P. van Dam, Berend Snel

**Affiliations:** Theoretical Biology and Bioinformatics, Department of Biology, Science faculty, Utrecht University, Utrecht, The Netherlands

## Abstract

Phylogenetic profiling in eukaryotes is of continued interest to study and predict the functional relationships between proteins. This interest is likely driven by the increased number of available diverse genomes and computational methods to infer orthologies. The evaluation of phylogenetic profiles has mainly focussed on reference genome selection in prokaryotes. However, it has been proven to be challenging to obtain high prediction accuracies in eukaryotes. As part of our recent comparison of orthology inference methods for eukaryotic genomes, we observed a surprisingly high performance for predicting interacting orthologous groups. This high performance, in turn, prompted the question of what factors influence the success of phylogenetic profiling when applied to eukaryotic genomes.

Here we analyse the effect of species, orthologous group and interactome selection on protein interaction prediction using phylogenetic profiles. We select species based on the diversity and quality of the genomes and compare this supervised selection with randomly generated genome subsets. We also analyse the effect on the performance of orthologous groups defined to be in the last eukaryotic common ancestor of eukaryotes to that of orthologous groups that are not. Finally, we consider the effects of reference interactome set filtering and reference interactome species.

In agreement with other studies, we find an effect of genome selection based on quality, less of an effect based on genome diversity, but a more notable effect based on the amount of information contained within the genomes. Most importantly, we find it is not merely selecting the correct genomes that is important for high prediction performance. Other choices in meta parameters such as orthologous group selection, the reference species of the interaction set, and the quality of the interaction set have a much larger impact on the performance when predicting protein interactions using phylogenetic profiles. These findings shed light on the differences in reported performance amongst phylogenetic profiles approaches, and reveal on a more fundamental level for which types of protein interactions this method has most promise when applied to eukaryotes.

## Introduction

The post-genomic era has provided us with a wealth of eukaryotic genomes of diverse and underrepresented phyla [1]. Most of the sequences in these new genomes are without precise function assignment and a challenge remains, protein function and interaction discovery [2,3]. Computational approaches that are available for large scale analyses of protein function and interactions include phylogenetic profiling. Phylogenetic profiling uses correlations of the presences and absences of groups of orthologous proteins (orthologous groups) across a set of species [4]. Phylogenetic profiling is a seemingly straightforward method proven to be a valuable alternative resource for studying functional relationships between proteins. Recently the method has played an integral part in identifying the cellular functional role of CENATAC that is a key player in a rare aneuploidy condition in humans [5], identifying eukaryotic reproduction genes [6], and identifying eukaryotic novel recombination repair genes [7].

The information in the phylogenetic profiles given by presence and absence patterns, are shaped by a diverse range of evolutionary forces. These forces include horizontal gene transfer, secondary endosymbiosis and gene loss. The method relies on the principle that proteins with a similar profile indicate that the proteins co-evolved due to them belonging to the same functional pathway or complex. There are countless observations that co-occurring proteins tend to interact [8–10]. Phylogenetic profiling can be a powerful tool for function prediction. By comparing, or even clustering, profiles of proteins with unknown function to those with known function enables us to infer to which complexes or functional pathways the uncharacterized proteins likely belong and, in turn, infer their function.

Multiple studies have shown the effectiveness of phylogenetic profiling in large scale analyses of eukaryotes [9,11], which has become possible with the large increase in genomic data and computational methods to (automatically) infer orthologies [12–14] or cluster genes [11]. However, benchmarking and analysing the performance of large-scale phylogenetic profiling has been limited to prokaryotes, for which good performance can be obtained when predicting protein interactions [15–18]. The performance decreases when benchmarking is done solely with eukaryotes or when eukaryotes are combined with prokaryotes [15,16]. Likely, the performance reduction is caused by the different forces driving eukaryotic genome evolution, compared to the dynamic pan genomes of prokaryotes where the interplay of rampant horizontal gene transfer of operons and loss of genes that create highly informative patterns.

We recently obtained a high protein interaction prediction performance in a large set of eukaryotes in the context of evaluating a diverse set of orthologous group inference methods [19]. The surprisingly high prediction performance only marginally depended on the orthologous group inference methods (which was the focus of the study), suggesting that its cause could be any of the other underlying choices. Therefore, a more elaborate analysis of the choices made for phylogenetic profiling is warranted. Here we evaluate in-depth the meta parameters influencing the performance of phylogenetic profiles in eukaryotes.

Multiple studies have understandably focused their analysis on reference genome selection or the amount of genomes/data needed to increase prediction performance [16–18,20,21]. Besides genome diversity and quality, we analyse orthologous groups and reference interactome selection. Our results demonstrate that an interplay of biological and technical aspects influence phylogenetic profiling. Most importantly, our results show that prediction performance is influenced not only by genome selection but mostly by orthologous and interactome selection.

## Results

Each results section describes the analyses of meta parameters encompassing five main concepts: genome quality, genome diversity, performance directed genome selection, orthologous group selection, and reference interactome selection. To rule out any orthology specific issues, we performed the analyses using two orthology inference methods, Sonicparanoid [13] and Broccoli [14]. Sonicparanoid performed the best in our previous study using phylogenetic profiles for protein interaction prediction [19]. We chose Sonicparanoid as the primary method, while broccoli serves to determine to what extent the results are contingent on a specific orthology method. The results for Broccoli can be found in Supplementary figures and are overall in agreement with the results of Sonicparanoid.

### 1. Lesser quality genomes have more effect on the prediction performance than higher-quality genomes

Phylogenetic profiles can be noisy due to multiple technical reasons, such as gene annotation and genome assembly errors. Consequently, the quality of genomes can be an essential factor, as profiles with a lot of noise would be akin to noisy gene expression or protein interaction measurements. We expect noisy genomes to give much weaker prediction performance. Given this expectation, the first meta parameter assessed was genome quality. We calculated genomes quality using two independent metrics, BUSCO [22] and one of our design (Supplementary figures and Methods and Materials). For clarity, we use only the BUSCO metric in the main text since both metrics generally agree with each other.

The BUSCO metric assesses genome completeness based on the (in our case) absence of single-copy orthologs that are highly conserved among eukaryotic species. The absences of these orthologs can result from incomplete draft genomes or false negatives in gene prediction, which in both cases leads to false absences of orthologs. We selected 50 high-quality genomes with the lowest BUSCO values, i.e., genomes with the least number of unexpected absences. We also selected 50 lower quality genomes with the highest BUSCO values, i.e., genomes with the most number unexpected absences (Fig 1.A.). We compared the quality filtered genome sets with 1000 randomly generated genome sets of 50 genomes each to see if quality-based selection differs from any random sampling of genomes.

The results show that the performance using the highest quality genomes with the least suspect absences falls within the distribution of random genome prediction performance (AUC: 0.765). In contrast, the lower quality genomes fall below the distribution of random prediction performance (AUC: 0.748) (inset Fig 1.B.). This suggests that it is more beneficial to filter out lesser-quality genomes than it is to select for high-quality genomes. This result is consistent between two independent scores of genome quality (S1 Fig).

With these results, it seems prudent to select genomes only based on quality when applying phylogenetic profiles. However, there is an inherent bias between genome quality and phylogenetic distribution (Fig 1.A). For instance, eukaryotes belonging to the Opisthokonta supergroup have overall lower BUSCO absences, biassing the selection of good genomes towards one eukaryotic supergroup. *A priori*, species diversity seems another meta parameter in genome selection with potential impact. In the next section, we will look at the diversity of species and how that influences phylogenetic profiling.

**Fig 1.**
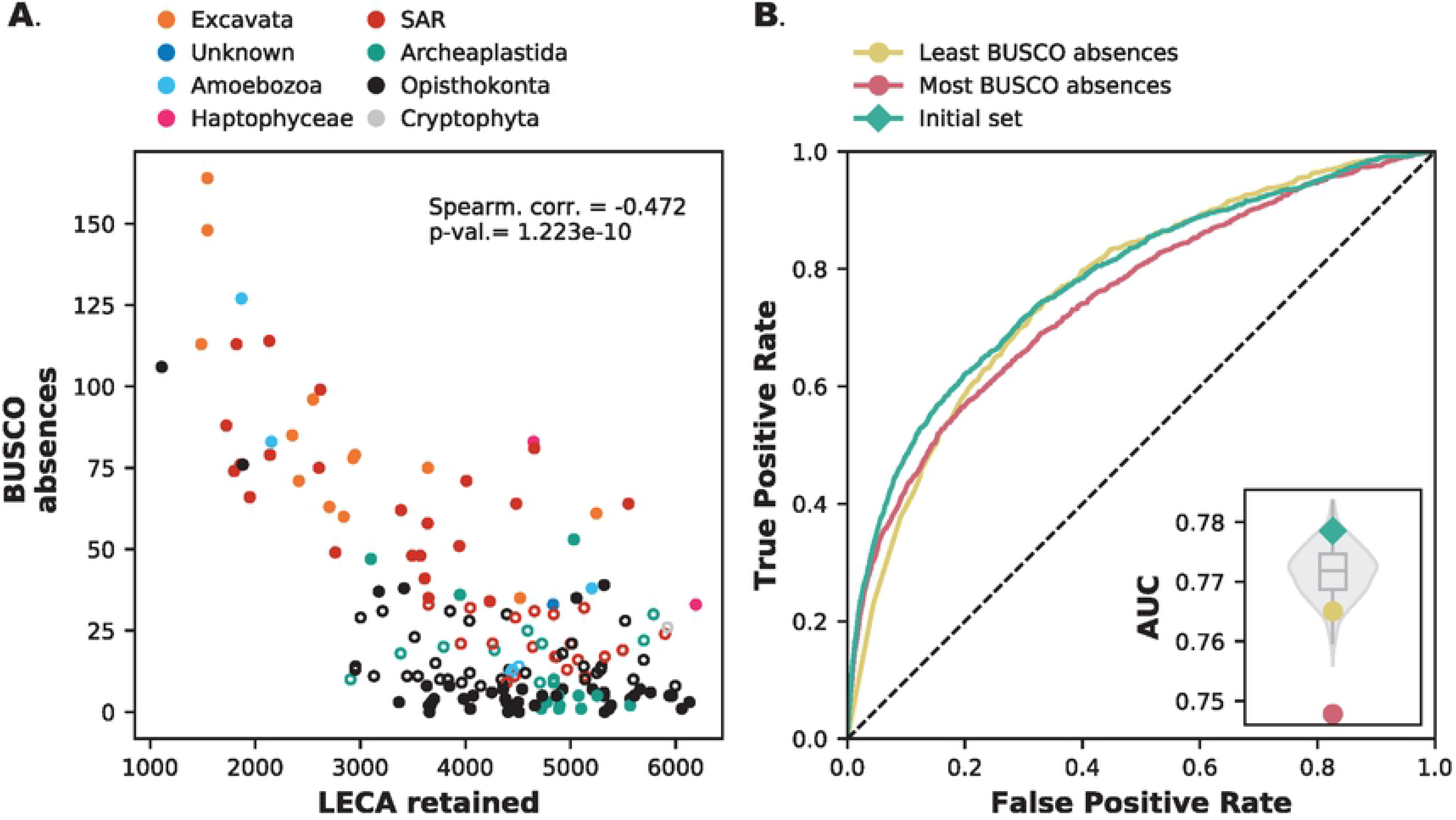
Lesser quality genomes have more impact on protein interaction prediction performance. **A.** BUSCO absences as a function of retained Last Eukaryotic Common Ancestor (LECA) orthologous groups in different species. Filled data points are the selected genomes for the prediction accuracy calculations. **B.** Receiver-operator Curve of two species sets (n = 50) with the most and least BUSCO absences. The inset gives the Area Under the Curve (AUC) values compared with the random backdrop of 1000 random species sets (violin plot) and the initial species set (teal diamond).

### 2. Genome diversity has little effect on prediction performance in eukaryotes

The diversity of species plays a role in the performance of phylogenetic profiles in prokaryotes [16]. We also expect high species diversity to improve how informative profiles are by giving high-resolution information on how genes co-evolve in different organisms. More species diversity allows to maximally discern the effect of evolutionary forces shaping co-evolving proteins, which might not be apparent in, e.g., an animal only data set. There will be no discernible and informative phylogenetic pattern in a homogeneous species set where most ancestral protein complexes are not frequently lost. A previous study showed that the maximum phylogenetic diversity in Bacteria gives the best predictive performance [18]. Here we want to test how maximal and minimal diversity affects prediction performance in eukaryotes.

We analysed the impact of eukaryotic diversity by selecting two sets of 50 genomes, one containing the most similar species (Fig 2.A.) and the other the most diverse species (Fig 2.B.) from our initial species set. The (dis)similarity was measured using an iterative all-vs-all comparison using the cosine distance between genomes and their orthologous group content. We started with the most diverse or similar species pairs and iteratively added to this set the species with the highest (dis)similarity until we obtained 50 genomes (Materials and Methods). We recalculated the protein-interaction prediction performance for both these sets. The prediction performance is lower than the initial set for both sets, but not worse than any randomly selected genome sets (AUC: 0.760 for the dissimilar set and AUC: 0.764 for the similar set) (Fig 2.C. inset).

**Fig 2.**
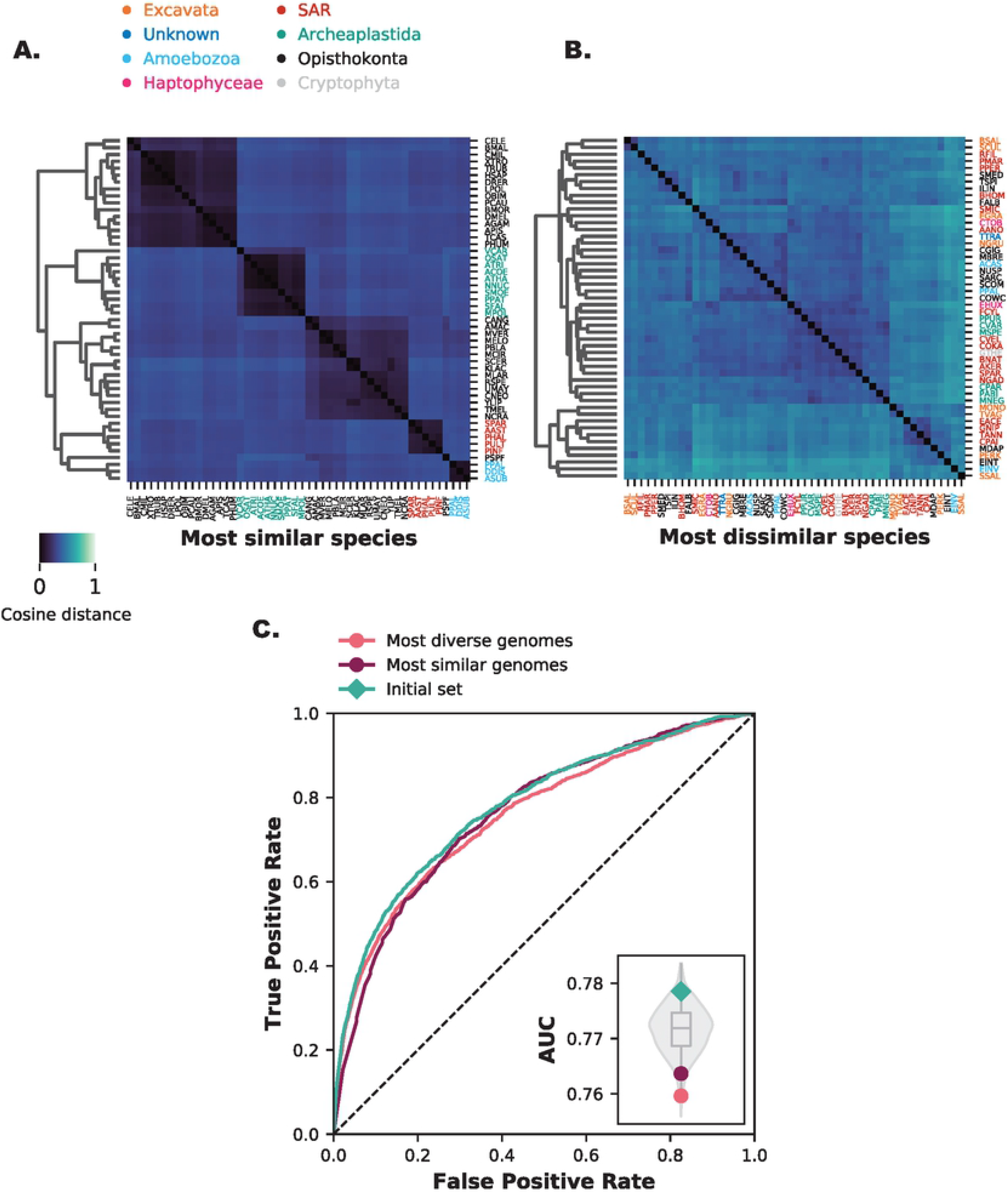
Both high and low diversity sets have little impact on protein interaction prediction performance. **A.** The most similar species form more clusters and are overall more similar to each other. **B.** The most diverse species show no clustering and are overall less similar to each other. **C.** Receiver-operator Curve of two species sets (n = 50) with the most diverse and most similar species. The inset gives the Area Under the Curve (AUC) values compared with the random backdrop of 1000 random species sets (violin plot) and the initial species set (green diamond).

Similar species will naturally show more cohesion in profiles, with little separation of protein co-evolution. Highly diverse species will naturally show more discordance, with little information left to see protein co-evolution. In both cases, there will not be a gene-specific signal. A combination of the two should give good separation of actual co-evolving genes. Together with the effects of genome quality, genome diversity can be an important factor for the performance of phylogenetic profiles. However, the interplay between these two factors is complex, and as we previously determined, high genome quality corresponds with lower phylogenetic diversity. To look into this further, we investigate the influence of single genomes on predictive performance.

### 3. Single influential genomes and their combined effect on prediction performance reveal the importance of the type of information in the profiles

Diversity and quality both impact performance and we expect it to have a combined influence on phylogenetic profiling. Instead of a priori selecting genomes based on a measure for each of these two criteria, we can also objectively evaluate the prediction performance by removing genomes from the initial species set one-by-one (Fig 3.A.). Genomes that decrease prediction performance when removed from the initial set we can consider as advantageous to phylogenetic profiling, while genomes that increase prediction performance when removed from the initial set we can consider as disadvantageous to phylogenetic profiling. We selected the top 50 advantageous and top 50 disadvantageous genomes to see whether these genomes together in their respective sets also influence the prediction performance.

For both the advantageous and disadvantageous set we can see a large difference in prediction performance (Fig 3.B.) and a larger difference than the selection based on the measures for either quality or diversity. With the advantageous genome set, the performance increases (AUC: 0.801). In contrast, for the disadvantageous set the performance drops (AUC: 0.730). Both values fall well outside the distribution of 1000 randomly generated genome subsets.

Although a large cumulative effect on performance because we used the genomes’ performance to select the genomes, it is still very interesting to see what these genomes share if it is not quality or diversity. We therefore examined the role of different genomes in these genome sets. Comparison of a large number of factors (S5 Fig) revealed that that the difference in prediction performance of the advantageous and disadvantageous genome sets is related to the (human) interactions retained in the genomes (Fig 3.C. and S5.A. Fig). The illogical absence ratios and the complete interactions present (or copresences) (Fig 3.C. I & II) show intermediate values for the disadvantageous genome set. At the same time, these values are either high or low for the advantageous genome set.

We can also directly relate the differences between these genome sets to how close the genomes in the sets are to the human genome. The cosine distance and the shared orthologous groups of the genomes with the human genome (Fig 3.C. III & IV) show intermediate values for the disadvantageous set, while the values are either high or low for the advantageous genome set. For the orthologous groups inferred by Broccoli this signal is even more pronounced (S5.B. Fig). A surprising finding is that the advantageous set contains numerous parasitic organisms (S1 Table).

In other words, genomes boosting the performance share either a lot or a little similarity with the reference interactome across a range of dimensions. Thus, phylogenetic profiling in eukaryotes benefits from genomes with a little or a lot of interactions present with regards to the reference interactome. These results reveal the importance of selecting genomes based on the evolutionary information contained within them relative to the query species, and is critical for high performance when predicting interacting proteins.

**Fig 3.**
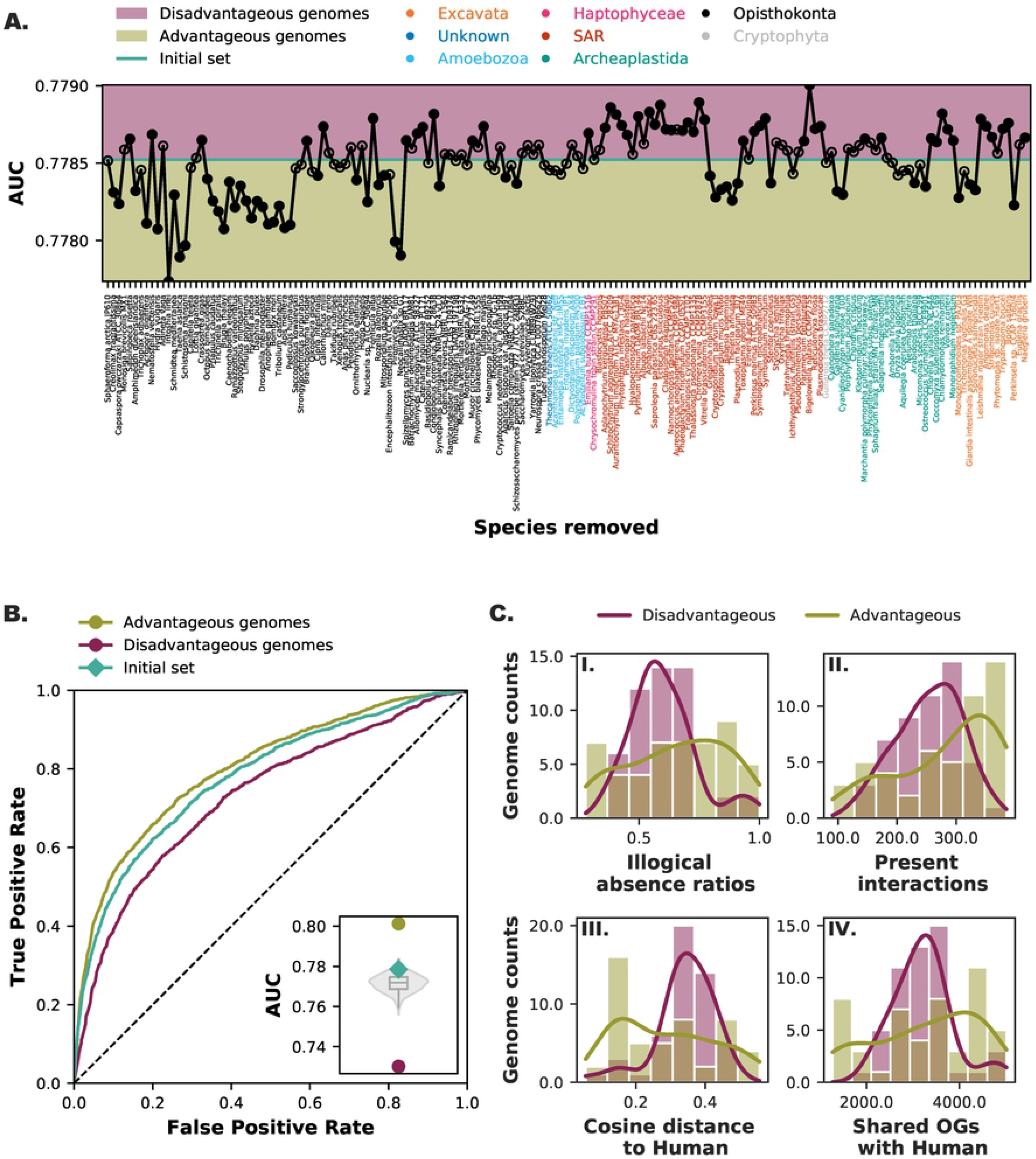
Influential genomes and their combined effect. **A.** Recalculated Area Under the Curve (AUC) values when a single species is removed from the initial species set. Genomes that increase the AUC value when removed can be considered disadvantages compared to the initial set when predicting protein interactions with phylogenetic profiles. Genomes that decrease the AUC value when removed can be considered advantageous for predicting protein interactions. Top 50 advantageous and top 50 disadvantageous genomes shown with the black fill in the scatter plot. **B.** Receiver-operator Curve of two species sets (n=50) with the most advantageous and disadvantageous genomes. The inset gives the Area Under the Curve (AUC) values compared with the random backdrop of 1000 random species sets (violin plot) and the initial species set (green diamond). **C.** Comparison of the counts (histogram) and kernel density estimates (line plot) of (I) illogical absence ratios (illogical absences divided by total interaction absences (co-absences + illogical absences)), (II) present interactions, (III) the cosine distance to human, and (IV) total shared orthologous groups with human.

### 4. Orthologous group (pre-)selection improves prediction performance by (inadvertently) enriching co-evolving proteins in profiles

Phylogenetic profiling benefits from clear modular co-evolution of proteins and subsets of proteins showing similar evolutionary behaviour [16,23]. A myriad of factors limit the modular co-evolution of interacting protein [24–26]. In previous research [19], which provides the starting meta parameters of this study, we evaluated orthology methods by their ability to recapitulate gene family dynamics in the Last Eukaryotic Common Ancestor (LECA). Consequently, the results so far are based on orthologous groups estimated to be in LECA. To see if this selection criterion was a factor in the strong performance, we performed phylogenetic profiling with other orthologous groups selections: groups estimated to be post-LECA, or groups not filtered on any criteria (post-LECA + LECA), i.e., the raw output of the orthology inference methods. We compared these orthologous group sets with 1000 subsets of randomly selected LECA orthologous groups. The prediction performance was indeed reduced (AUC: 0.691 post-LECA and AUC: 0.734 all orthologous groups) compared to that of LECA orthologous groups or any randomly selected set of orthologous groups (Fig 4.). After some reflection, a myriad of explanations likely factor into this effect. Profiles of LECA proteins have many losses, and thus a lot of information (entropy) (S6 and S8 Figs). Profiles of post-LECA proteins have less loss and, by definition, are restricted to specific lineages, and thus contain less information. Combining LECA and post-LECA orthologous groups produce a set of phylogenetic profiles with an overall much lower similarity.

**Fig 4.**
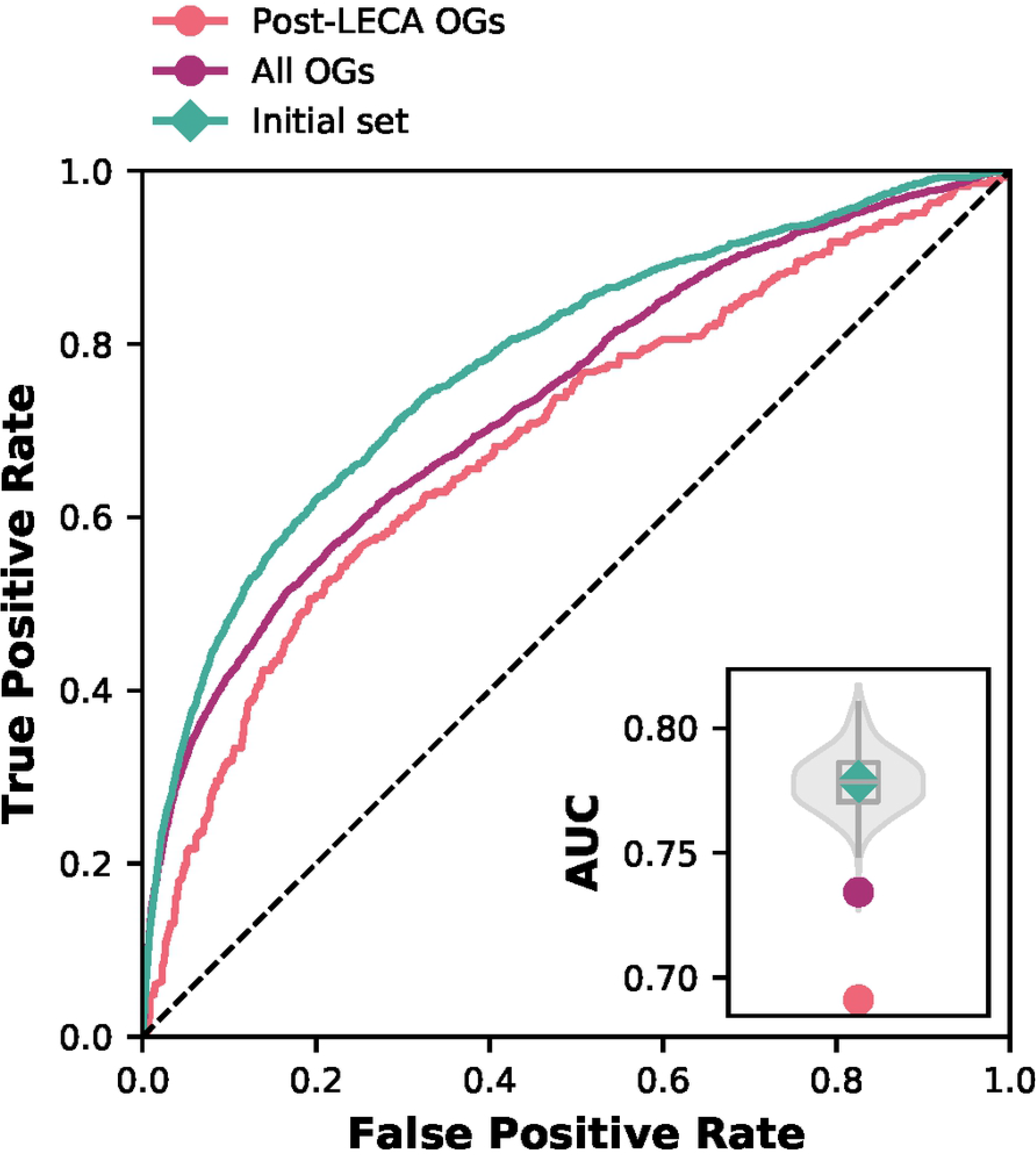
Orthologous group selection has a large impact on prediction performance. Receiver-operator Curve of post-LECA orthologous groups and unfiltered orthologous groups. The inset gives the Area Under the Curve (AUC) values compared with the random backdrop of randomly selected LECA OGs (violin plot) and the initial species set (green diamond).

We now have identified a key meta parameter choice explaining why our previous research found such high performance. However, it is unclear what the reason for this effect is and why for specific pairs of proteins, one protein was in LECA and the other not. This separation could be biological reality, i.e., innovations in the evolution from LECA to human, or issues in orthology assignment, i.e., one protein is evolving much more rapidly that causes the protein’s predicted orthologous group to give an artifactually lineage-specific distribution in the profile. Consequently, the protein is falsely inferred as a more recent addition or innovation. Manual inspection of this set (S9 Fig) does not obviously point towards one of the explanations. It is likely a combination of factors, including orthology prediction errors (e.g., oversplitting) and actual lineage-specific additions/inventions. In any case, the meta parameter of orthologous group selection is perhaps easily overlooked or made implicitly in the OG creation itself. Still, it is highly impactful, and our results show that OG selection improves prediction performance by enriching co-evolving proteins in profiles.

### 5. Choice of reference interactome and interaction filtering improves prediction performance by increasing the amount of co-evolving proteins and quality of interactions

Phylogenetic profiling attempts to predict which pairs of proteins are part of the same function, pathway, or complex. The performance of phylogenetic profiles can be measured using a data set of proteins that interact or are otherwise functionally linked. For example, we can take KEGG pathways as measuring units, as done in the STRING database [27]. However, these pathways often have an excess of 30 proteins and not all of them are expected to have the mutual functional dependence that results in co-evolution. This unwantedly biases the predictor by having supposedly interacting proteins with little correlation. Similarly, if we would take a very small well-curated set of compact/short linear metabolic pathways as was used to seed the CLIME searches [11], then the choice of what protein pairs to count as false negatives becomes difficult. Hence, our decision in previous work was to parse human interactions from BioGRID to contain only interactions found in at least five independent studies (Methods and Materials). This filtering of interactions has been repeatedly demonstrated to effectively increase the quality and reduce the noise in the interaction data [27,28]. Moreover, the same data set contains a very good indication of which proteins are not functionally related. Proteins that are well studied and repeatedly surface in high throughput assays and are subject to repeated investigations are indeed likely to have no functional relation since these proteins are evidently never identified to interact.

The results in section 3 (Fig 3.C.) reinforce the notion that the reference interaction set plays a role in the performance of predicting interacting proteins. For these reasons, we analysed how the filtering and choice of reference interactome influences protein interaction prediction performance in eukaryotes. Using an unfiltered human protein interaction dataset reduces the prediction performance from an AUC of 0.779 to an AUC of 0.638 (Fig 5.A.). This performance is also lower than any set of randomly selected LECA orthologous groups (inset). The quality of the interaction data used clearly plays a role in prediction performance, i.e., if we take a noisy “ground truth” it turns out to be difficult to predict this truth. It is difficult to predict interactions with a set littered with false, virtually random, pairs.

**Fig 5.**
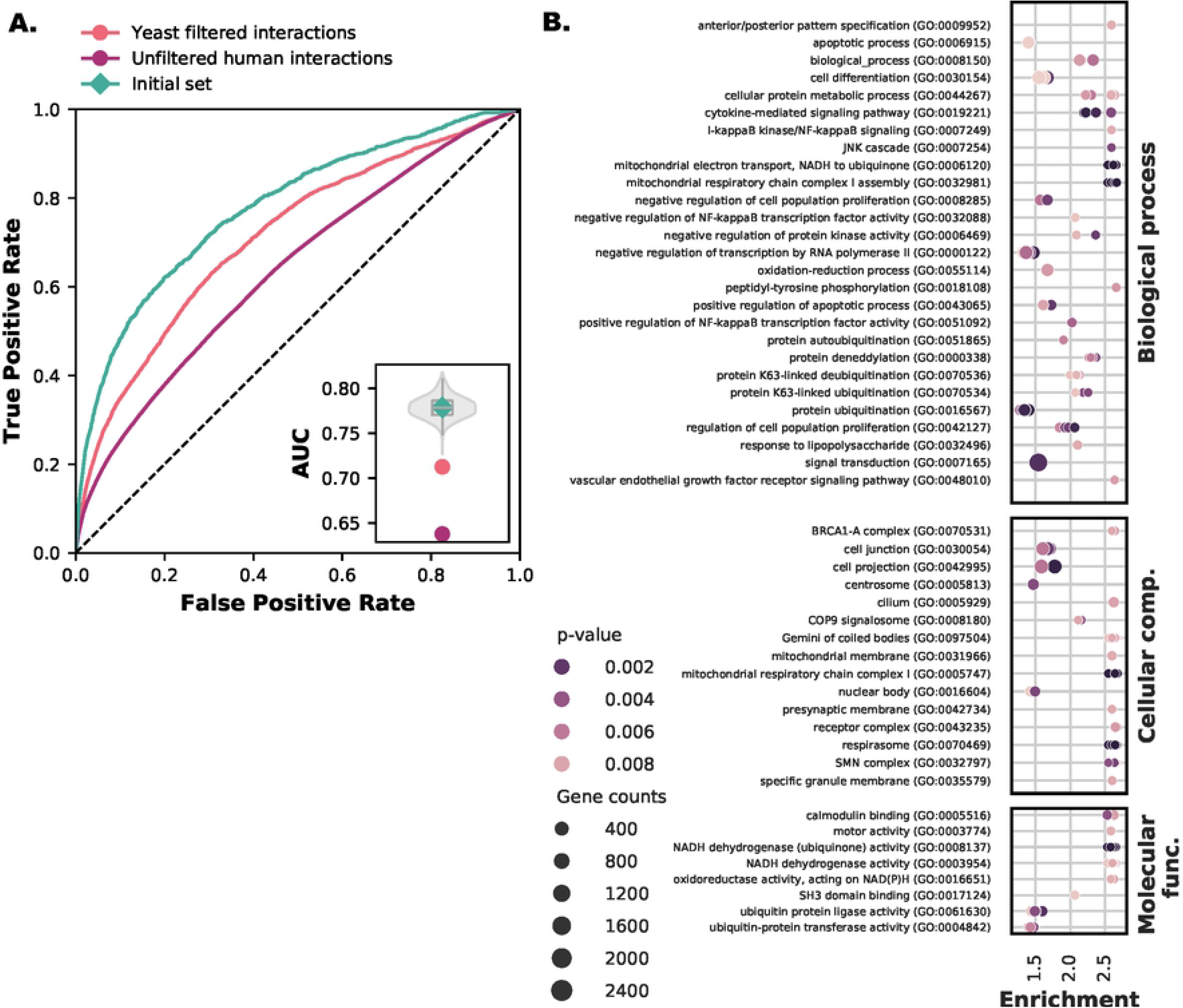
Interactome selection is important for prediction performance. **A.** Receiver-operator Curve of post-LECA orthologous groups and unfiltered orthologous groups. The inset gives the Area Under the Curve (AUC) values compared with the random backdrop of randomly selected LECA orthologous groups (violin plot) and the initial species set (green diamond). **B.** GO-enrichment analysis for genes enriched in interactions present in only human compared to interactions present in human and yeast. Orthologous groups can contain multiple genes. We randomly selected genes from an orthologous group to generate a new sample and population sets ten times and recalculated the enrichment (shown by multiple points in the figure rows).

We further analysed the choice of reference organism for protein interactions. Specifically for eukaryotes, the prediction performance was sensitive to the reference species for protein interactions [21]. Yeast has been the organism of choice as the reference interaction set for eukaryotes. Yeast is a popular model organism that has been extensively researched, and it is with yeast that many protein-protein interaction high throughput methods were pioneered. As a result, we also expect the interaction data of yeast to be of higher quality than that of human and, consequently, interaction predictions to be better.

We used *Saccharomyces cerevisiae* interactions from BioGRID (Materials and Methods) filtered with the same number of publications strictness criterion. Surprisingly, and contrary to for instance [6], the human interaction set performed better with an AUC of 0.779 compared to the yeast interaction set with an AUC of 0.713 (Fig 5.A.). One reason could be that ascomycete fungi and yeast in particular, has lost many co-evolving LECA complexes found in most eukaryotes [29,30]. These losses include Complex I, essential functions in chromatin modification [31], spliceosomal introns and RNAi machinery giving patchy patterns of canonical Dicer and Argonaute [32], ciliary genes [8,33], and the WASH complex [9,11,34]. These observations prompted us to look at the GO term enrichment of interacting LECA orthologous groups that contain only human genes versus interacting LECA orthologous groups that have both human and yeast genes.

We indeed find evidence of multiple genes belonging to ancestral complexes enriched in the human interaction set (Fig 5.B.), including enrichment in more straightforward GO terms related to mitochondria and respiration (e.g., GO:0005747, GO:0006120, GO:0032981 and GO:0070469), cilium (e.g., GO:0005929) and spliceosomal components (e.g., SMN complex GO:0032797). We also find evidence in higher-level GO terms that at lower levels reflect complexes known to be present in human and absent in yeast (S2 Table), such as chromatin modification (e.g., GO:0042127). For the broccoli inferred orthologous groups, more enriched GO terms reflect at lower-level complexes known to be present in human and not in yeast: Argonaute and Dicer (GO:0010629, GO:0048471 and GO:0030426), WASH (GO:0005814, GO:0005856 and GO:0043005) and chromatin modification (e.g., GO:0007399) (S10 Fig and S3 Table).

Even though there are more yeast than human interactions present in multiple species, the entropies of the profiles participating in yeast interactions are lower (S11 Fig). This observation and the GO analysis reveal a clear reason why the performance we reported is high relative to others. Namely, we use the human reference interaction set with ancestral complexes that have been frequently lost throughout eukaryotic evolution and are absent in yeast.

## Discussion

Phylogenetic profiling is complicated due to many biological and technical issues. These issues include the complex histories of proteins and the choices in the meta parameters for phylogenetic profiling, such as the quantity, quality and diversity of reference genomes and annotations. Meta parameter choices in phylogenetic profiling has been extensively studied in mostly prokaryotes, where generally the focus is on the choice of reference genomes, phylogenetic profile methods, and/or the amount of data. We focus on eukaryotes and investigate qualitative different meta parameters for phylogenetic profiling. We showed that phylogenetic profile performance when predicting protein interactions is influenced by a complex interplay of multiple technical and biological parameters.

Genome diversity plays an important role in prediction performance for prokaryotes [16,18]. In contrast, our measures of eukaryotic diversity did not significantly influence prediction performance. Selecting lesser-quality genomes has a larger effect on prediction performance, while selecting higher-quality genomes does not. Genome selection and the interplay between quality and diversity does matter. However, other meta parameters have a much larger impact on prediction performance, such as the amount of information in the phylogenetic profiles in relation to the reference interaction dataset. This discrepancy suggests that more complex feature selection procedures should be explored for reference genome set selection, especially since (non-linear) interactions between subsets of genomes and combinations of subsets could drastically boost performance.

Other meta parameters, such as orthologous group and reference interactions, have a much larger effect than genome selection. Some results make a lot of sense from a technical point of view. For example, low quality/noisy functional data (unfiltered BIOGRID) or mixing phylogenetic profiles that are at least 50% inconsistent (post-LECA + LECA orthologous groups) have poor performance. A drawback to filtering out post-LECA orthologous groups is that we remove lineage-specific interactions that are still a part of a protein complex and show clear co-evolution. Our analysis shows that we should consider these often hidden choices when encountering large differences in performance between reported studies.

One very counterintuitive finding is that the yeast interaction set showed lower predictive performance. Compared to the human interaction set, yeast should be of equal quality by all accounts, arguably even better. Together with the observation that LECA orthologous groups performed better than post-LECA orthologous groups, this suggests that the performance of phylogenetic profiles in eukaryotes is optimal for modules that fulfil a very particular set of conditions. These modules (i) were present in LECA, (ii) were repeatedly lost in eukaryotic lineages, and (iii) the genes in the module conserved most of their function. This observation fits with notable examples from the WASH complex and cilium [8,9,11], or proteins with great success in predicting its components like the minor spliceosome [5] and RNAi machinery genes including Dicer and Argonaute [32]. These biological patterns should explain the very strong signal found by studies such as [9,11]. Note, both studies show very strong signals for complexes as well as pathways, which we excluded due to the problem of defining a quality negative interaction set.

In conclusion, we find that for eukaryotes more genomes and better-quality genomes are not necessarily better. It is instead the type of information in the genomes. The information in these genomes is not directly related to larger genomes, for instance parasites increase prediction performance. Instead, the information is related to the interactions of the reference species present in a given genome. Genome selection has a minor influence compared to orthologous groups selection and interactome selection, which both greatly improve the performance when predicting protein interactions. Interactome and orthologous group selection is likely the major source for the large variance in reported performances. Ancestral complexes that are repeatedly lost are responsible for the strong performance of phylogenetic profiles in eukaryotes and it is these hidden choices in orthologous group selection that we should consider when we find large differences in performance between studies.

## Material and Methods

### 1. Initial datasets and methods

We started our investigation from the analysis done in our previous work [19], to investigate the influence of different parameters on the performance of predicting protein-protein interactions using phylogenetic profiles. We showed a relatively high prediction performance using a large set of diverse eukaryotes and orthologous groups inferred to be in the Last Eukaryotic Common Ancestor (LECA). This reference set is called the initial set. Any changes that we made are changes in this initial set. In the sections below, we will briefly describe the composition of this initial set and the methods we used to obtain it.

#### 1. 1. Large scale eukaryotic dataset and LECA orthologous groups

We inferred orthologous groups on a diverse genome set of 167 eukaryotes using different orthologous group inference methods in our previous work. For this analysis, we chose the best performing method regarding protein interaction prediction, Sonicparanoid (version 1.3.0) [13]. To rule out any large orthology specific issues during our current analyses, we chose at least one other method: Broccoli (version 1.0) [14].

Ancestral eukaryotic complexes have been lost together multiple times [35]. Phylogenetic profiles should benefit from this clear modular evolution of proteins. Therefore, we selected orthologous groups estimated to be in LECA. Briefly, we inferred LECA orthologous groups using the Dollo parsimony approach [36] with additional strict inclusion criteria [19]. The Dollo parsimony method assumes that genes can be gained only once while minimizing gene loss. Before we assigned an orthologous group to LECA, it must be in at least three supergroups (See Supp. Table X) distributed over the Amorphae and Diaphoretickes (previously known as opimoda and diphoda) [37].

#### 1. 2. Phylogenetic profiling and measuring co-occurrence of proteins

We constructed phylogenetic profiles by determining the presence (1) and absence (0) of orthologous groups in 167 species. To evaluate prediction accuracy, we obtained a higher quality reference interaction set by filtering the human BioGRID interaction database (version 3.5.172 May 2019) [38,39]. BioGRID contains physical interactions between proteins. We filtered this interaction set to keep non redundant interaction pairs found in at least five independent studies (PubMed ID’s). The number of independent studies is a measure of how thoroughly these proteins were investigated and how receptive the proteins are to high-throughput measurements. We mapped the interacting genes to their corresponding orthologous groups.

We used the best performing negative protein interaction set from our previous analyses [19]. We inferred this negative set by taking pairs of interacting proteins that were found to be interacting at least five times, but not with each other. This excludes the possibility that the negative set contains interacting proteins that were not found due to manifold technical reasons.

To calculate the (dis)similarities between phylogenetic profiles we used the from our previous analysis best performing distance measure, the cosine distance.

### 2. Genome selection procedures

We compared the results of all the genome selection procedures to 1000 sets of genomes randomly selected to exemplify that the differences in prediction accuracies are not due to random variations in genome composition. We calculated the protein interaction prediction performance for each of these random genome sets.

#### 2. 1. Selecting better and worse quality genomes

To measure the quality of the genomes, we used two quality metrics. The first metric is the out of the box BUSCO metric that works by calculating the absences of highly conserved single-copy orthologs [22]. The BUSCO Eukaryota database (odb9) was aligned to the genomes using the hmmsearch alignment tool from the HMMER package 3.1b2 (dated February 2015) [40]. We took the HMM specific quality score given by BUSCO to validate the hits in the alignments.

The second metric is of our own devising. The second quality metric we used was the Illogical absences (IA) metric of our design. We added this second independent metric to remove the dependence of quality on a single measure to establish the completeness of the genomes and gene prediction. The IA metric calculates the number of absences of protein interaction partners, which we termed Illogical absences. Illogical absences follow from the assumption underlying phylogenetic profiling that interacting proteins are often evolutionary conserved. Therefore, it can be considered suspect when a protein interaction partner is absent. A possible reason could be that the absence is due to gene prediction, genome annotation or even homology detection errors.

We selected the strongest interacting orthologous group pairs by selecting their phylogenetic profiles with the least cosine distance. This selection removes the complex interplay between interacting groups of orthologs. For every interacting orthologous group pair, we calculated the absences of interaction partners in every species. These absences we termed illogical absences or the IA metric.

We can consider the genomes with the most BUSCO absences and illogical absences as lesser quality genomes. In contrast, we can consider the genomes with the least BUSCO absences and illogical absences as higher-quality genomes. We selected 50 genomes of lesser-quality and 50 genomes of higher-quality for each of the metrics and recalculated the protein-protein interaction prediction performance.

#### 2. 2. Selecting highly diverse and similar genomes

We calculated the pairwise cosine distance between all species with the presence and absence profiles of LECA orthologous groups to obtain species sets of maximum diversity and maximum similarity. We then iterated through the resulting pairwise distance matrix and selected the maximally distant pairs for the diverse set or minimally distant pairs for the similar set. Before adding a species of a species pair to a set, we checked to see if the species also had a distance above a certain arbitrary threshold to the other species in the growing set (cosine value ≥ 0.38 for the dissimilar, cosine value ≤ 0.58 for the similar set). We did this until we obtained the desired amount of 50 genomes per set. The maximum diverse and maximum similar genome sets were each used to recalculate the protein-protein interaction prediction performance.

#### 2.3. Selecting single influential genomes and their combined effect on prediction performance

We removed genomes one-by-one from the initial species set of 167 eukaryotes to see how the different genomes influence the performance of protein interaction prediction with phylogenetic profiling. We recalculated the performance for each of these 167 sets. The 50 genomes that increased the performance compared to the initial species set the most when removed from the initial set were labelled as disadvantageous. The 50 genomes that decreased the performance the most when removed from the initial set were labelled advantageous. For both the disadvantage and advantageous set we recalculated the protein interaction prediction performance.

### 3. Gene and interactome selection procedures

We compared the results of the orthologous group selection procedures to randomly selected LECA orthologous groups to exemplify that the differences in prediction accuracy is not due to random variations in orthologous group composition. We made a thousand LECA orthologous group sets containing a random selection of 63% of the orthologous groups. We calculated each of these set’s protein interaction prediction performance.

#### 3. 1. Selecting orthologous groups

In our initial species set, we used orthologous groups estimated to be in LECA (Methods section 1.1.). We took the raw output of the orthology inference methods and filtered out the LECA orthologous groups to get a set that contains post-LECA orthologous groups. We also recalculated the prediction performance with the raw output of the orthology prediction methods, which is all inferred orthologous groups.

#### 3.2. Selecting different reference interactomes

We compared the five PubMed ID filtered human BioGRID set with the unfiltered human BioGRID dataset. Every interaction with less than five pubIDs is now included as well. Removing the five PubMedID filter should indicate how quality filtering of reference interactions influences prediction performance.

We selected next to the human interactions the *Saccharomyces cerevisiae* BioGRID interaction database (version 3.5.175 July 2019) [39] to analyse the influence of the reference interactome. We filtered the interactions to keep only the interaction pairs found in at least five publications (PubMed ID’s). We followed the same procedure as with the human interaction set (Methods section 1.2.).

Following this analysis, we hypothesized that the drop in prediction performance for yeast is caused by the loss of ancestral protein complexes in yeast. To test this, we chose interacting LECA orthologous groups that contained only human genes (sample set) and calculated the enrichment to the set with interacting LECA orthologous groups containing human and yeast genes (population set). We calculated the enrichment using the following equation: 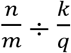 where n is the total number of genes associated with a GO term (Downloaded GO terms Januari 2021 biomart) in the sample set (overlap), m is the total numbers of genes in the sample set, k is the total number of genes associated with a specific GO term in the population set, and q is the total number of genes in the population set. Since enrichment does not work well for small overlaps, we filtered for a minimum overlap (n) of 3. Enrichment was considered significant for p-values below 0.01. Since orthologous groups can contain multiple genes, we randomly selected genes from an orthologous group to generate a new sample and population sets ten times and recalculated the enrichment.

## Supporting information

**S1 Fig. Illogical absences and genome quality selection based on illogical absences using Sonicparanoid inferred orthologous groups (OGs). A.** Illogical absences as a function of retained LECA OGs in different species. We see that where most Opisthokonta scores similarly low with the BUSCO metric, they score lower with the IA metric indicating a difference between the two metrics. However, the performance of genomes selected with both metrics are similar to each other. Filled data points are the selected genomes for the prediction accuracy calculations. **B.** Receiver-operator Curve of two species sets (n = 50) with the most and least illogical absences. The inset gives the Area Under the Curve (AUC) values compared with the random backdrop of 1000 random species sets (violin plot) and the initial species set (teal diamond). Human has a perfect score of 0 illogical absences since the interactions are from the human reference interactome. Therefore, we did not select human for the genome set.

**S2 Fig. Lesser quality genomes have more impact on protein interaction prediction performance also for Broccoli inferred orthologous groups (OGs). A.** BUSCO and Illogical absences as a function of retained LECA OGs in different species. Filled data points are the selected genomes for the prediction accuracy calculations. **B.** Receiver-operator Curve of two species sets (n = 50) with the most and least BUSCO and illogical absences. The inset gives the Area Under the Curve (AUC) values compared with the random backdrop of 1000 random species sets (violin plot) and the initial species set (teal diamond).

**S3 Fig. Both high and low diversity sets have little impact on protein interaction prediction performance also for Broccoli inferred orthologous groups. A.** The most similar species form more clusters and are overall more similar to each other. **B.** The most diverse species show no clustering and are overall less similar to each other. **C.** Receiver-operator Curve of two species sets (n = 50) with the most diverse and most similar species. The inset gives the Area Under the Curve (AUC) values compared with the random backdrop of 1000 random species sets (violin plot) and the initial species set (green diamond).

**S4 Fig. Influential genomes and their combined effect for Broccoli inferred OGs. A.** Recalculated Area Under the Curve (AUC) values when a single species is removed from the initial species set. Genomes that increase the AUC value when removed can be considered disadvantages compared to the initial set when predicting protein interactions with phylogenetic profiles. Genomes that decrease the AUC value when removed can be considered advantageous for predicting protein interactions. Top 50 advantageous and top 50 disadvantageous genomes shown with the black fill in the scatter plot. **B.** Receiver-operator Curve of two species sets (n=50) with the most advantageous and disadvantageous genomes. The inset gives the Area Under the Curve (AUC) values compared with the random backdrop of 1000 random species sets (violin plot) and the initial species set (green diamond). **C.** Comparison of the counts (histogram) and kernel density estimates (line plot) of (I) illogical absence ratios (illogical absences divided by total interaction absences (co-absences + illogical absences)), (II) present interactions, (III) the cosine distance to human, and (IV) total shared orthologous groups with human.

**S5 Fig. Correlations between multiple parameters in the advantageous and disadvantageous genome set.** Given for **A.** Sonicparanoid and **B.** Broccoli inferred orthologous groups (OGs). From top to bottom (or left to right) the interactions that are co-absent; illogically absent; and present; the ratio of illogical absences to total absences; number of OGs shared with the human genome; the cosine distance to the human genome; LECA OGs loss (Dollo parsimony inferred); species (lineage) specific loss; (clade) ancestral loss; and the difference in AUC from the initial set AUC when a genome is removed.

**S6 Fig. Entropy of phylogenetic profiles that have interactions.** Given for **A.** Sonicparanoid and **B.** Broccoli inferred orthologous groups (OGs). From top to bottom, the entropy is shown in profiles for LECA, post-LECA and all OGs. Median entropy is presented with a black arrow. Mann-Whitney U test shows significant difference between distributions of LECA, post-LECA and all OGs, p-value < 0.001.

**S7 Fig. Orthologous group selection has a large impact on prediction performance also for Broccoli inferred orthologous groups (OGs).** Receiver-operator Curve of post-LECA orthologous groups and unfiltered orthologous groups. The inset gives the Area Under the Curve (AUC) values compared with the random backdrop of randomly selected LECA OGs (violin plot) and the initial species set (green diamond).

**S8 Fig. Dollo parsimony inferred loss of LECA and post-LECA orthologous groups (OGs).** Given for **A.** Sonicparanoid and **B.** Broccoli inferred OGs. Mann-Whitney U test shows significant difference between distributions.

**S9 Fig. Groups of interacting orthologous groups (OGs) where one is in LECA (always the last row in a group subplot) and the others are not.** The profiles are sorted according to the species tree.

**S10 Fig. Interactome selection is important for prediction performance. A.** Receiver-operator Curve of post-LECA and unfiltered orthologous groups (OGs) of Broccoli. The inset gives the Area Under the Curve (AUC) values compared with the random backdrop of randomly selected LECA OGs (violin plot) and the initial species set (green diamond). **B.** GO-enrichment analysis for genes enriched in interactions present in only human vs. interactions present in human and yeast. OGs can contain multiple genes. We randomly selected genes from an OG to generate new sample and population sets 10 times and recalculated the enrichment (shown by multiple points in the figure rows).

**S11 Fig. Interactions of human and yeast interactome present in different species (left) and entropy for LECA profiles that have interactions in human and yeast (right).** Given for **A.** Sonicparanoid inferred and **B.** Broccoli inferred orthologous groups (OGs). Median values are presented with the arrows. Mann-Whitney U test shows significant difference between distributions.

**S1 Table. Species table for species used in this study.** Green marked species are the species that are in the advantageous set, and red marked species in the disadvantageous set (Sonicparanoid). The measured values are shown in S5 Fig.

**S2 Table. GO-enrichment table for Sonicparanoid inferred orthologous groups (OGs).** Since there can be multiple genes in an OG, we randomly selected one of the genes for the GO-enrichment analysis. We did this ten times, creating ten foreground and background sets (set_num). These values are shown in Fig 5.

**S3 Table. GO-enrichment table for Broccoli inferred orthologous groups (OGs).** Since there can be multiple genes in an OG, we randomly selected one of the genes for the GO-enrichment analysis. We did this ten times, creating ten foreground and background sets (set_num). These values are shown in S10 Fig.

